# Attention capture outside the oculomotor range

**DOI:** 10.1101/2020.08.12.249003

**Authors:** Nina M. Hanning, Heiner Deubel

## Abstract

Neurophysiological studies demonstrated that attentional orienting is performed by fronto-parietal brain areas which also play an important role in oculomotor control. Accordingly, several studies claimed that exogenous attention can only be allocated to where we can potentially make an eye movement, i.e. within the oculomotor range. We tested this assumption by assessing the disruptive effect of a salient distractor at locations within and beyond participants’ oculomotor range. Participants rotated their heads ~38° leftwards to prevent them from performing large rightward saccades. In this posture, participants fixated the screen center and focused their attention on a location on the left side of the screen, where they had to discriminate the orientation of a visual noise patch. While assessing visual orientation sensitivity – an established proxy of visual attention – at this endogenously attended location, we flashed a salient cue either at the attended location or at various locations inside or outside their oculomotor range. We found that whenever the salient cue occurred at a location other than the endogenously attended location, it withdrew visual attention and significantly disrupted endogenous attentional orienting. Crucially, this effect occurred regardless of whether the cue was presented within or beyond participants’ oculomotor range, demonstrating that exogenous events equally grab our attention both inside and outside the eyes reach. Since spatial exogenous attention was attracted unrestrictedly toward locations to which no saccade could be executed, the coupling of attention and eye movement control is less tight than, for example, the prominent “Premotor Theory of Attention” suggests. Rather, attention can be shifted freely over the entire visual range, independent of pathological and physiological limitations of the eye movement system.

## MAIN TEXT

Neurophysiological studies have demonstrated that attentional orienting is associated with activity in fronto-parietal brain areas that play a pivotal role in oculomotor control, such as the lateral intraparietal cortex (LIP), the frontal eye fields (FEF), and the superior colliculus (SC) [e.g., 1]. Accordingly, based on the influential *premotor theory of attention,* which posits that even *covert* shifts of spatial attention in the absence of eye movements are elicited by preceding activation in the oculomotor system [2], it has been claimed that attention can only be allocated to where we can potentially make an eye movement [3]. There are two forms of covert spatial attention: *Exogenous attention* is automatic, stimulus-driven, and transiently deployed in ~100 ms. Conversely, *endogenous attention* is voluntary, goal-driven, and deployed in a slower (~300 ms) and sustained manner [4]. Notably, it has been postulated that only exogenous attention, but not endogenous attention, would be restricted to locations within the so-called *oculomotor range* that is accessible by saccadic eye movements [5,6]. To test this claim, we used a dissociation approach that allowed us to evaluate exogenous attention shifts to locations within and beyond observers’ oculomotor range via their disruptive, attention capturing costs for endogenous attention. We found that salient events equally grab exogenous attention both inside *and* outside the oculomotor range, demonstrating that exogenous attention can shift to locations not reachable by the eyes.

Across two experiments, 7 observers rotated their heads ~32° leftwards (**Figure 1A**), which prevented them from performing rightward saccades larger than ~8° (degrees of visual angle). The required head rotation angle was individually determined prior to the experiment and monitored with an electromagnetic motion tracking device (see **Figure S1** for individual head rotation angles). We used a two-alternative forced-choice discrimination task (**Figure 1B**) based on oriented pink noise patches [7] (**Figure 1C**) to assess covert spatial attention across four locations on the horizontal meridian (±6° and ±10° relative to central eye fixation). Due to the leftward head rotation, the right distal patch (+10°) lied beyond the oculomotor range (i.e., it could not be reached by the eyes but was still visible). Observers were instructed to attend to the left distal location (−10°; within the oculomotor range), as the brief orientation signal (50 ms; tilted clockwise or counterclockwise) they had to discriminate was most likely to occur there. In Experiment 1, the discrimination signal occurred in 75% of trials at the −10° location (and in 25% of trials at one of the three remaining locations with equal probably); in Experiment 2 the signal occurred at the −10° location in 100% of trials. While observers aimed to maintain endogenous attention at the −10° location, we flashed a salient 50 ms cue at one of the four locations (−10°, −6°, +6°, +10°; with equal probability). As a highly salient visual event, this cue can be assumed to strongly attract exogenous attention (typically within ~100 ms [4]). By evaluating observers’ visual sensitivity – an established proxy for visual attention – at the endogenously attended (−10°) location 100 ms following cue presentation, we inferred the attention-capturing effect of the exogenous cue depending on whether it occurred inside or outside the oculomotor range. While we expected high sensitivity when the exogenous cue matched the endogenously attended location (−10°), sensitivity at the endogenously attended location should be decreased when the cue attracted exogenous attention away from it. Critically, if the cue attracted exogenous attention even at +10° outside the oculomotor range, this would demonstrate that exogenous attention can shift to locations not reachable by the eyes – falsifying previous claims [3,5,6].

**Figure 1.**
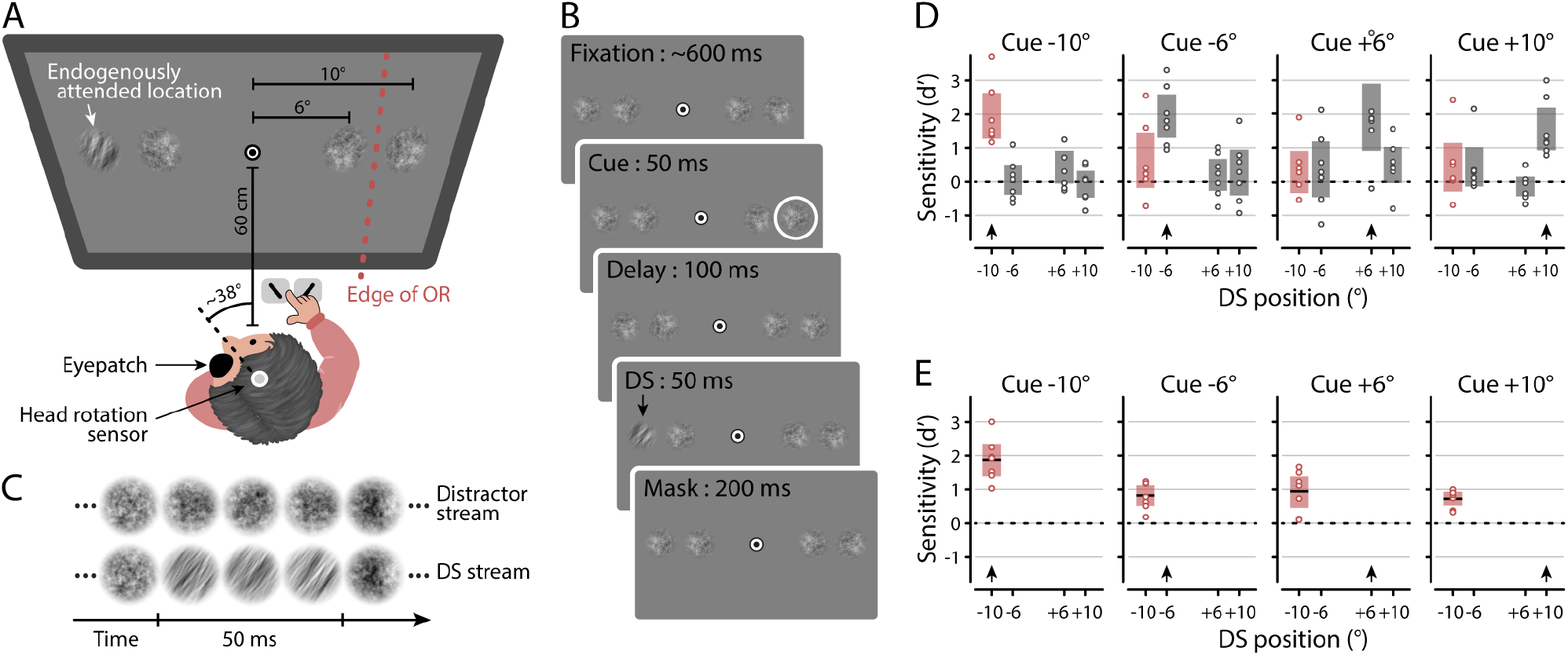
Attention capture task design and results. (A) Setup. Observers viewed the stimuli with their left eye patched and their head rotated leftwards. Four pink noise streams were presented at four locations, covering two eccentricities (±6° and ±10°) from a central fixation target. The +10° patch fell outside observers’ oculomotor range (OR). (B) Each pink noise stream consisted of a succession of randomly generated pink noise patches that flickered at 60 Hz. The discrimination signal (DS) stream included a sequence of orientation-filtered pink noise patches (50 ms), showing a 40° clockwise or counterclockwise tilt relative to the vertical axis. (C) Experimental task. After a fixation period, an exogenous, circular white cue was flashed around one randomly chosen noise stream. Observers were instructed to ignore this cue, maintain central eye fixation, and report the orientation of the DS presented 100 ms after the cue (**Movie S1**). (D and E) Visual sensitivity in Experiment 1 (D) and Experiment 2 (E) at the endogenously attended −10° location (red whisker plots), and the other locations (−6°, +6°, +10°; blue whisker plots) as a function of cue location (indicated by the black arrow at the bottom of each plot). Horizontal lines within each whisker plot indicate the mean visual sensitivity (d′) at the respective test position. Error bars depict 95% confidence intervals, dots represent individual observer data, dashed lines mark chance level.

For both Experiment 1 (**Figure 1D**) and Experiment 2 (**Figure 1E**), we evaluated visual sensitivity (*d’*) at the endogenously attended −10° location depending on whether the exogenous cue matched or did not match this location. We found that visual sensitivity at the endogenously attended −10° location was significantly higher when the cue occurred at the same location (**Cue −10°**; Experiment 1: 2.08 [1.72, 2.45] – mean [95% CI]; Experiment 2: 1.89 [1.65, 2.13]), as compared to when it occurred at a neighboring location within the oculomotor range (**Cue −6°** Experiment 1: 0.70 [0.27, 1.13], *p* < 0.001; Experiment 2: 0.84 [0.69, 0.99], *p* < 0.001; **Cue +6°** Experiment 1: 0.34 [0.01, 0.67], *p* < 0.001; Experiment 2: 0.95 [0.71, 1.19], *p* < 0.001). Thus, as expected, the cue decreased visual performance at the endogenously attended location by exogenously attracting attention away from it. Crucially, and in direct contradiction to the predictions of previous work [5,6], this disruptive effect also occurred when the cue was presented at +10°, well outside the oculomotor range – as indicated by an equally reduced sensitivity at the endogenously attended −10° location (**Cue +10°** Experiment 1: 0.49 [0.11, 0.88], *p* < 0.001; Experiment 2: 0.76 [0.66, 0.86], *p* < 0.001).

The findings reported here demonstrate that exogenous events equally grab our attention both inside and outside the oculomotor range, and are in line with recent evidence from our lab showing concordant attentional benefits beyond the reach of saccadic eye movements [8]. The automatic shift of attention to a distracting cue outside the oculomotor range observed here is an exogenous effect that cannot be explained by voluntary, endogenous orienting. As spatial exogenous attention was attracted unrestrictedly toward locations to which no saccade could be executed, the present findings question the coupling of exogenous attention and eye movement control proposed by the premotor theory of attention [2] and its variants [3,5,6]. Our results are in line with a recent study demonstrating that, unlike previously claimed [9], pathological oculomotor restrictions are not necessarily associated with corresponding attentional deficits [10]. Instead, exogenous attention can be shifted freely over the entire visual range, independent of limitations imposed by the eye movement system.

## SUPPLEMENTAL INFORMATION

Supplemental information contains acknowledgements, author contribution statement, a detailed description of the experimental procedure, a Figure S1 depicting the head rotation data, and a Movie S1 demonstrating the trial sequence.

## REFERENCES

1. Jonikaitis, D., & Moore, T. (2019). The interdependence of attention, working memory and gaze control: behavior and neural circuitry. Current Opinion in Psychology, 29, 126–134.

2. Rizzolatti, G., Riggio, L., Dascola, I., &Umiltá, C. (1987). Reorienting attention across the horizontal and vertical meridians: evidence in favor of a premotor theory of attention. Neuropsychologia, 25(1), 31–40.

3. Smith, D. T., & Schenk, T. (2012). The premotor theory of attention: time to move on? Neuropsychologia, 50(6), 1104–1114.

4. Carrasco, M. (2011). Visual attention: The past 25 years. Vision Research, 51(13), 1484–1525.

5. Smith, D. T., Schenk, T., & Rorden, C. (2012). Saccade preparation is required for exogenous attention but not endogenous attention or IOR. Journal of Experimental Psychology: Human Perception and Performance, 38(6), 1438.

6. Casteau, S., & Smith, D. T. (2020). Covert attention beyond the range of eye-movements: Evidence for a dissociation between exogenous and endogenous orienting. Cortex, 122, 170–186.

7. Hanning, N. M., Deubel, H., & Szinte, M. (2019). Sensitivity measures of visuospatial attention. Journal of Vision, 19(12), 1–13.

8. Hanning, N. M., Szinte, M., & Deubel, H. (2019). Visual attention is not limited to the oculomotor range. Proceedings of the National Academy of Sciences, 116(19), 9665–9670.

9. Rafal, R. D., Posner, M. I., Friedman, J. H., Inhoff, A. W., & Bernstein, E. (1988). Orienting of visual attention in progressive supranuclear palsy. Brain, 111(2), 267–280.

10. Masson, N., Andres, M., Carneiro Pereira, S., Pesenti, M., & Vannuscorps, G. (in press). Typical exogenous covert shift of attention without the ability to plan eye movements. Current Biology.

