## Supplemental Information for "Attention capture outside the oculomotor range"

### Acknowledgements

We would like to thank Daniela Gresch for her valuable support in data collection and Marisa Carrasco and Marc Himmelberg for helpful comments and discussions. This research was supported by Deutsche Forschungsgemeinschaft (DFG grant DE336/6-1 to H.D.).

### Author Contributions

N.M.H. conceived and designed the experiments, performed research, and analyzed the data; N.M.H. and H.D. wrote the paper.

### Declaration of Interests

The authors declare no conflict of interest.

### Supplemental Experimental Procedure

**Observers.** Sample sizes were determined based on previous work [S1]. 7 observers (3 females, ages 23–31 years-old, one author) participated in Experiment 1, 7 observers participated in Experiment 2 (3 females, ages 23–33 years-old, one author, 4 of whom also participated in Experiment 1). All observers had normal vision and except for one author (N.M.H.) were naive as to the purpose of the experiment. The protocols for the study were approved by the ethical review board of the Faculty of Psychology and Education of the Ludwig-Maximilians-Universität München and conducted in accordance with the Declaration of Helsinki. All observers provided written informed consent.

**Apparatus.** Head rotation was recorded via a sensor of a Polhemus Liberty 240/8 electromagnetic motion tracking device (Polhemus Inc., Colcherster, Vermont, USA) and a personal toolbox. The sensor was attached on top of observers' head using an EEG-cap (position CZ). The exact amount of head rotation was determined to ensure that each observer's oculomotor range ended in between the right proximal (+6°) and the distal (+10°) location. Eye position of the dominant right eye was recorded using an EyeLink 1000 Tower Mount eye tracker (SR Research, Osgoode, Ontario, Canada) at a sampling rate of 1 kHz. After initial head rotation and placement, the eye tracker was calibrated (and recalibrated whenever the head was moved). Calibration targets were presented within 8° from fixation, such that they fell within observers' oculomotor range. According to SR Research, the eye tracker used allows precise tracking of the eye position within the range required for our paradigm (up to ~12° on the horizontal axis). Manual responses were recorded via a standard keyboard. The experimental software controlling display, response collection, and eye and head tracking was implemented using Matlab (MathWorks, Natick, MA, USA), using the Psychophysics [S2, S3] and EyeLink toolboxes [S4], running on a Dell Precision T1500 Intel Core i5 computer (Round Rock, Texas, USA). Stimuli were presented at a viewing distance of 60 cm on a 21-in SONY GDM-F500R CRT screen (Tokyo, Japan) with a spatial resolution of 1024 by 768 pixels and a vertical refresh rate of 120 Hz.

**Experimental design.** Observers sat in a dimly illuminated room, with their left eye patched and their head rotated about 32° to the left (see Main Fig. 1), positioned on a chin rest. The head rotation angle was determined individually prior to the experiment (to ensure that the +10° location fell outside each observer's oculomotor range; see **Figure S1** for individual head rotation angles) and monitored with an electromagnetic motion tracking device. Each trial began with observers fixating a central fixation target (FT) comprising a black (~0 cd/m2) and white (~120 cd/m2) bull's eye (radius 0.25°) on a gray background (~60 cd/m2). Once stable fixation was detected within a 2.0° radius virtual circle centered on FT for at least 200 ms, four pink (1/f) noise streams [S5] (radius 1.5°) appeared on the horizontal axis at -10°, -6°, +6° and +10° relative to FT (positive values correspond to the right side of the screen). Each noise stream consisted of randomly generated pink noise patches (mean luminance ~60 cd/m2) windowed by a symmetrical raised cosine (radius 1.5°, sigma 0.5°), refreshing at 60 Hz (Fig. 1B). After a random fixation period between 400 and 800 ms, a cue (white annulus, radius 1.5°, ~120 cd/m2) was flashed for 50 ms around one randomly selected noise stream (note that cues flashed at +10° fell outside the oculomotor range due to the head rotation). Observers were instructed to ignore the task-irrelevant cue and continuously keep fixation at the FT. 100 ms after cue onset the discrimination signal (DS) was presented. The DS consisted of an orientation-filtered noise stimulus, displaying a rotated pattern, tilted 40° clockwise or counterclockwise relative to the vertical. In Experiment 1, observers were informed that the DS in 75% of trials would appear within the noise stream at the -10° location (and in 25% trials randomly within one of the noise streams at the -6°, +6°, or +10° location). In Experiment 2, observers were informed that the DS appeared at the -10° location in all trials. After 50 ms the DS was masked by the reappearance of non-oriented noise. 200 ms after DS onset the noise streams disappeared and observers reported the orientation of the DS via button press. They were informed that in neither experiment the cue was predictive of the DS location, and that their orientation report was non-speeded (they were instructed to take their time to rest and blink before initiating the next trial by giving their response). Observers received auditory negative feedback for incorrect responses.

Observers performed at least 720 trials in Experiment 1 (8 experimental blocks) and at least 140 trials in Experiment 2 (1 experimental block) and took breaks after each block to ensure the maximum comfort despite the unusual head position. Eye fixation was monitored throughout each trial, and observers received auditory feedback whenever their eye position left a radius of 2.0° around the FT. Incorrect trials were aborted and repeated in random order at the end of each block.

A threshold task preceded the main experiments to ensure a consistent level of discrimination difficulty across observers. The threshold task was identical to the main experiments but observers did not rotate their head, the cue occurred with equal likelihood at any of the 4 locations, and observers were informed that the DS would always be presented at the cued location. We used a procedure of constant stimuli and randomly selected the orientation filter strength (corresponding to the visibility of the orientation tilt) out of 5 linear steps of filter widths. By fitting cumulative Gaussian functions to the discrimination performance data gathered in this threshold task, we determined the filter width corresponding to 80% correct discrimination performance and used this value in the respective main experiment.

**Eye data pre-processing.** We scanned the recorded eye-position data offline and only included trials in which no blink occurred and correct eye fixation was maintained within a 2.0° radius centered on FT throughout the trial. In total we included 4940 trials in the analysis of the behavioral results of Experiment 1 (on average 705.71 ± 34.29 trials per observer) and 977 trials of Experiment 2 (on average 139.57 ± 0.79 trials per observer).

**Behavioral data analysis.** We compute observers' sensitivity in the orientation discrimination task, *d'* = z(hit rate) - z(false alarm rate) separately for each location, depending on the cued location. We defined hits as "clockwise" responses to a clockwise orientation, and false alarms as "clockwise" responses to a counterclockwise stimulus. If the observed proportion correct was equal to 100% or 0%, we substituted these performance levels with 99% and 1%, respectively. Performances below chance level (d' = 0 corresponding to 50%) were transformed to negative d' values.

Whisker plots to visualize the data show single observer sensitivity (represented by dots) that we averaged across observers (represented by black lines), as well as the 95% confidence interval (indicated by colored bars). For all statistical comparisons we resampled our data and derived *p*-values by locating any observed difference on the permutation distribution (difference in means based on 1000 permutation resamples). All reported differences remained significant after Bonferroni multiple-comparison correction. All files are available from the OSF database URL: https://osf.io/u8vsx.

### Supplemental Figure

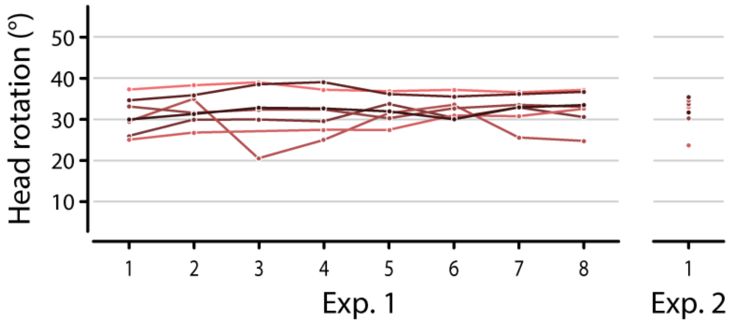


**Figure S1**. Individual observers rotation average head rotation angles over the time course of Experiment 1 (Left; eight blocks) and Experiment 2 (Right; one block). Different colors show different observers.The average rotation angle was 32.20 ± 1.48° (mean ± SEM) in Experiment 1 and 31.68 ± 1.61° in Experiment 2. Note that the head rotation of one observer was not saved for one block. Excluding this particular block does not change our behavioral findings.

### Supplemental Movie

**Movie S1.** Trial demonstration. Rotate your head approximately 32° to the left and fixated the central fixation "bull's eye" with your left eye closed. Start the video while keeping eyes steady. Try to ignore the circular white cue and discriminate the tilt of the oriented noise patch presented at the outer left location. (Cue presented at +10°; orientation (clockwise) presented at -10°).

### Supplemental References

S1 Hanning, N. M., Szinte, M., & Deubel, H. (2019). Visual attention is not limited to the oculomotor range. *Proceedings of the National Academy of Sciences*, *116*(19), 9665-9670.

S2 Brainard, D. H. (1997). The psychophysics toolbox. *Spatial Vision*, *10*(4), 433-436.

S3 Pelli, D. G. (1997). The VideoToolbox software for visual psychophysics: Transforming numbers into movies. *Spatial Vision*, *10*(4), 437-442.

S4 Cornelissen, F. W., Peters, E. M., & Palmer, J. (2002). The Eyelink Toolbox: eye tracking with MATLAB and the Psychophysics Toolbox. *Behavior Research Methods, Instruments, & Computers*, *34*(4), 613-617.

S5 Hanning, N. M., Deubel, H., & Szinte, M. (2019). Sensitivity measures of visuospatial attention. *Journal of Vision*, *19*(12), 1-13.
